# Augmentation of self-motion perception with synthetic auditory cues

**DOI:** 10.1101/2025.07.14.664677

**Authors:** Roie Karni, Adam Zaidel

## Abstract

People who suffer from vestibular loss or damage have difficulty maintaining balance and perceiving their own motion in space (self-motion). Sensory augmentation of vestibular information, via other senses, could improve these functions. Here, we tested whether synthetic auditory signals carrying self-motion information can be integrated with vestibular cues. Twenty healthy participants experienced self-motion stimuli in a 3D motion simulator, comprising vestibular (inertial motion), and/or synthetic auditory “motion” cues. The auditory cues, presented via stereo headphones, comprised a series of beeps, with motion speed encoded by beep rate, and heading direction (in the horizontal plane) encoded by simulating the sound to emanate from that direction. In each trial, participants experienced a single-interval self-motion stimulus (vestibular, auditory or combined), and their task was to discriminate its heading direction (two-alternative forced-choice, right/left of straight ahead). A slight heading conflict (Δ = ±6°) was introduced in the combined-cue condition to measure empirical cue weights. Combined auditory-vestibular thresholds were significantly lower (improved) compared to vestibular alone (p < 0.001). But integration was suboptimal – vestibular cues were overweighted, and combined-cue thresholds were larger than predicted by Bayesian-optimal weighting. Interestingly, combined-cue thresholds were better predicted by the empirically observed weights. Thus, humans can integrate synthetic auditory cues with natural vestibular cues to improve self-motion perception. However, synthetic cues are underweighted. This suggests that weighting is determined not only by cue reliability but also by perceived relevance, with (unnatural) synthetic cues potentially considered less relevant to the task. Training might be required to achieve reliability-based integration of synthetic and natural cues.

## Introduction

Individuals who suffer from vestibular loss or damage have difficulty perceiving their own motion in space (self-motion perception) and maintaining balance (Iwasaki & Yamasoba, 2015). However, impaired vestibular function is not limited to disease or pathology. Rather, it is a normal aspect of natural aging (Allen et al., 2016; Anson & Jeka, 2016). This puts older people at high risk of falling (Iwasaki & Yamasoba, 2015; Jacobson et al., 2008), which is the leading cause of hospital admission and accidental death in the elderly (Bergen et al., 2016; Florence et al., 2018). Improving vestibular function for people with vestibular damage and for the elderly can improve stability, reduce the risk of falls, and thereby enhance their quality of life and physical well-being.

One approach to replace or augment lost or damaged vestibular function is by directly stimulating the vestibular nerve internally (similar to a cochlear implant; Golub et al., 2014; Guinand et al., 2015). However, this solution involves surgery, which is invasive and costly. Also, it may be technically challenging to selectively stimulate specific nerve fibers related to different motion directions and types (e.g., linear vs. rotational; Sluydts et al., 2019) to attain a naturalistic and coherent experience. Therefore, it is also important to also explore non-invasive alternatives.

Sensory substitution (or augmentation) is a method to replace (or augment) sensory information lost from one sense via a different sense. It has many advantages: it is cheap, non-invasive, and can be engineered to provide high-resolution and specific signals (Bach-y-Rita et al., 1969; Maidenbaum et al., 2014). Thus, sensory substitution may offer a viable solution for impaired vestibular function. In fact, the vestibular sense may be particularly suitable for this, as it is inherently multisensory – there is no ‘primary’ vestibular cortex; rather, vestibular inputs project to a wide range of multisensory cortical areas (Lopez et al., 2012; Rancz et al., 2015). Moreover, the vestibular system demonstrates remarkable cross-modal plasticity (Lopez et al., 2012; Rancz et al., 2015; Shalom-Sperber et al., 2022; Zaidel, 2024; Zaidel et al., 2011, 2013; Zeng et al., 2023). Therefore, the vestibular sense may be an ideal candidate for sensory substitution or augmentation.

Several studies have investigated sensory substitution or augmentation of vestibular function. In these studies, vestibular information was provided using vibrotactile cues (Mahmud et al., 2022; Sienko et al., 2008; Velázquez, 2010), electrical stimulation of the tongue (Bach-y-Rita et al., 2005), auditory cues (Dozza et al., 2005; Hasegawa et al., 2017; Hegeman et al., 2005), or multi-modal cues (Fortin et al., 2008; Huffman et al., 2010). The results show that receiving vestibular information via other senses can improve function. However, the solutions proposed in these studies generally focus on improving one specific functional aspect: maintaining an upright posture in order to prevent falls. Accordingly, they typically monitor the tilt of an individual’s trunk (deviation from upright posture) and warn when the tilt is too large. But solutions focused on tilt have inherent limitations: i) a person does not always want to avoid tilt; rather, tilt is part of normal function (e.g., leaning forward when getting up from a chair, or bending down). ii) Tilt-based solutions do not capture the full spectrum of vestibular functionality. The vestibular system provides rich information beyond tilt, including rotation (e.g., yaw rotation around the vertical axis) and linear acceleration. Therefore, broader aspects of sensory substitution and augmentation of vestibular functions beyond tilt require investigation.

Auditory cues may provide a good substrate for vestibular sensory substitution or augmentation. Firstly, auditory cues from the environment improve balance (Gandemer et al., 2017; Hasegawa et al., 2017; Negahban et al., 2017; Vitkovic et al., 2016), and patients with vestibular deficits exploit these auditory cues to a greater degree (Shayman et al., 2018; Stevens et al., 2017; Vitkovic et al., 2016). This suggests that individuals extract spatial information from auditory cues to support balance and perhaps also self-motion perception. Additionally, auditory cues contain a rich set of features, including pitch, duration, loudness, timbre, and spatial location (Burton, 2015). These multiple features can be used to encode different aspects of self-motion simultaneously by combining them into a composite auditory signal. Indeed, composite auditory cues comprising different combinations of auditory features have been successfully used to augment other senses, most notably vision (Auvray et al., 2007; Maidenbaum et al., 2014; Spagnol et al., 2017).

In this study, we investigated sensory augmentation of vestibular signals, with a focus on linear self-motions in the horizontal plane (as a first step toward augmenting vestibular signals more generally). An advantage of focusing first on linear self-motion perception is that some vestibular implant solutions are designed to specifically stimulate rotational self-motions, i.e., to replace semi-circular canal function (Boutros et al., 2019; Chow et al., 2021). Thus, addressing linear cues (noninvasively) can fill an important gap, and complement such vestibular implants. For this purpose, we developed a synthetic auditory cue that encodes linear velocity and heading direction in the horizontal plane. Participants experienced vestibular, auditory, or combined (vestibular-auditory) self-motion stimuli in a motion simulator and performed a task of heading discrimination. We investigated the following questions: do synthetic auditory cues improve heading discrimination? Is combined-cue performance in line with Bayesian-optimal integration? And if not, what model of cue-combination well-describes the observed data?

We found that performance with combined-cue (vestibular-auditory) stimuli was significantly better than with vestibular stimuli alone. Thus, synthetic auditory cues indeed improve heading discrimination. However, combined-cue performance was not in line with Bayesian-optimal integration: (a) combined-cue thresholds exceeded Bayesian predictions, and (b) vestibular cues were overweighted compared to Bayesian predictions. Lastly, we found that combined-cue thresholds were better predicted when using the empirically observed cue weights, extracted from the data, rather than Bayesian-predicted weights.

## Methods

### Participants

Twenty adults (9 women and 11 men; mean age ± SD = 27.80 ± 6.06 years, age range: 19 - 45), recruited from Bar-Ilan University, participated in this study. The study was approved by the Internal Review Board at Bar-Ilan University. All participants signed informed consent prior to partaking in the study and received monetary compensation for participation.

### Experimental setup and stimuli

The participants were seated comfortably in a car seat, which was mounted on a motion platform (MB-E-6DOF/12/1000, Moog Inc.), and restrained safely in the seat with a 4-point harness (Fig. 1A). The participant’s head was supported by a head support with lateral extensions to limit head movement (Black Bear, Matrix Seating Ltd.). A central fixation point was presented throughout the duration of the experiment using a head-mounted display (HMD; Oculus Rift) worn by the participants. Because minor head movements were possible, participants were instructed to keep their heads straight and still, and to focus on the fixation point, throughout the experiment.

**Figure 1.**
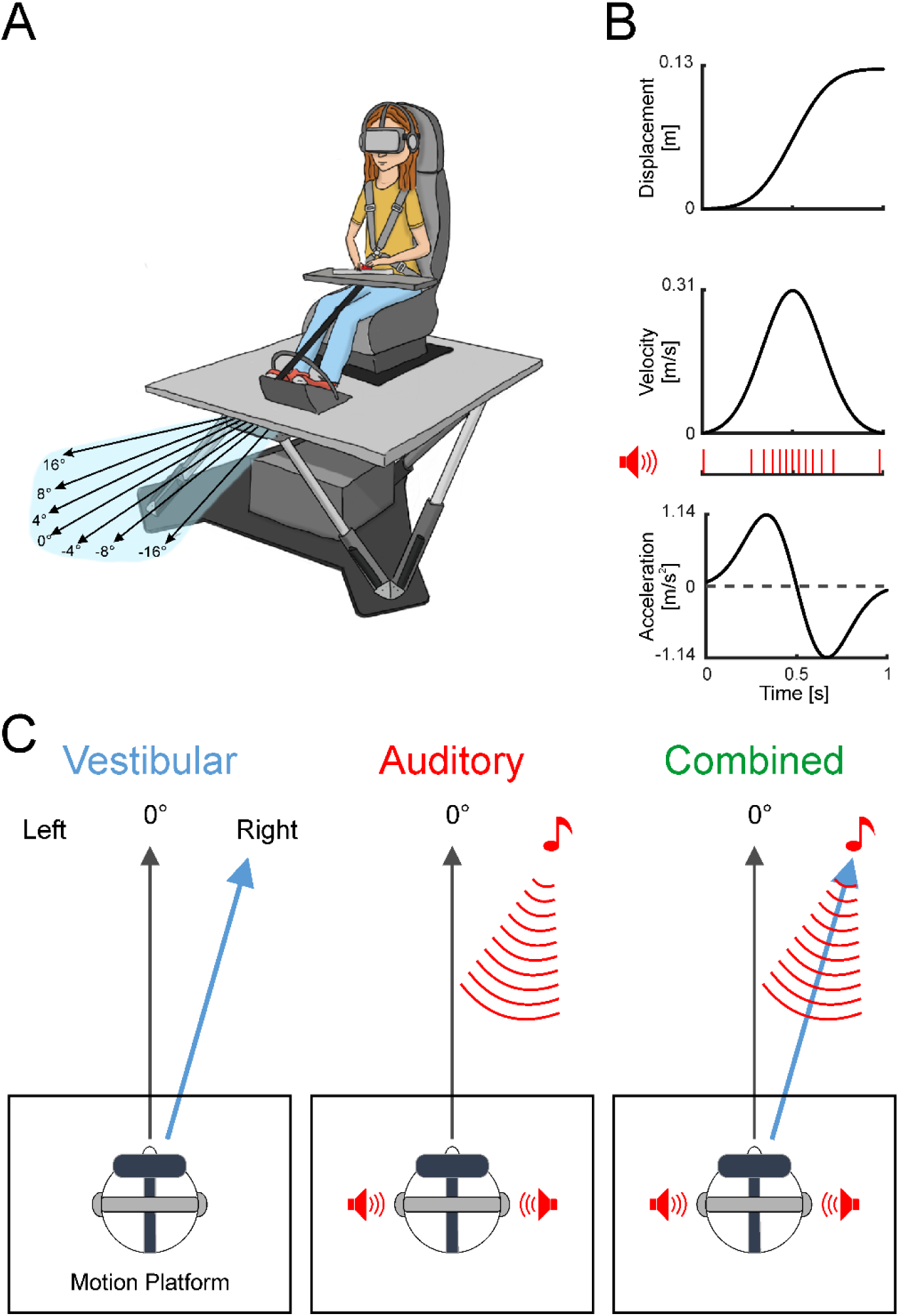
Experiment setup and stimuli. (A) Schematic of the multisensory motion simulator. Black arrows on the light blue background depict several different self-motion headings (in the horizontal plane). (B) Motion profile of the self-motion stimuli. The red bars under the velocity motion profile indicate the timing of the auditory beeps. (C) Schematics (viewed from above) of the three types of self-motion stimuli: vestibular, auditory, and combined vestibular-auditory (left, center, and right, respectively). The blue arrow depicts an example vestibular stimulus heading, rightward of center. The red arcs depict a synthetic auditory ‘heading’ stimulus (designed to emanate from the corresponding heading, played via stereo headphones). The combined stimulus comprises both vestibular and auditory stimuli, presented together. The black arrows mark ‘straight-ahead’ (0°).

The vestibular self-motion stimuli were similar to those used in previous studies (Yakubovich et al., 2020; Zaidel et al., 2011, 2015). The stimuli comprised single-interval linear motions in a primarily forward-moving direction, with slight deviations to the right or left of straight ahead. Possible stimulus heading values were distributed logarithmically around straight ahead: ±16°, ±8°, ±4°, ±2°, ±1°, ±0.5°, or ±0.25°, where zero represents straight ahead, and positive (or negative) values represent headings to the right (or left) of straight ahead. The stimuli followed a Gaussian velocity motion profile with 1 s duration and total displacement 0.13 m (peak velocity 0.31 m/s and peak acceleration 1.14 m/s^2^; Fig. 1B). Although additional (e.g., somatosensory) cues might also be present during inertial motion, we refer to this condition as ‘vestibular’ because performance strongly relies on intact vestibular labyrinths (Gu et al., 2007).

We designed a synthetic auditory stimulus to encode velocity and heading direction in accordance with the vestibular self-motion stimuli. This was done in two steps: first, a canonical signal representing the stimulus velocity was created. Velocity was encoded by beep rate (1 beep/cm traveled). The velocity profile in all trials was the same (irrespective of heading), hence, the number of beeps and their timing were identical across trials (Fig. 1B, red bars). Second, the heading direction was encoded by making the auditory signal sound like it was emanating from a source in that particular direction. Specifically, a binaural cue was synthesized using a 3D head-related transfer function (HRTF; Matlab audio toolbox) to sound like it was coming from a point in space one meter away from the participant. The location relative to the participant remained fixed throughout the stimulus; hence it provided an egocentric directional cue, and not an allocentric locational one. The auditory stimuli were played through noise-canceling headphones (Bose QC30) worn by the participant throughout the experiment. They were well above the detection threshold, but not uncomfortably loud.

To test and quantify multisensory cue weighting, a slight discrepancy (Δ) was introduced between the vestibular and auditory headings when presented in combination. By convention, Δ > 0 indicates that the vestibular and auditory headings were offset to the right and left, respectively, each by Δ/2 (vice versa for Δ < 0). For this study, we used Δ = ±6°. This was presumed to be within the range of integration (rather than segregation) based on previous work that showed visual-vestibular integration for |Δ| ≲ 10° (Acerbi et al., 2018; De Winkel et al., 2017) and typical vestibular thresholds.

### Heading discrimination task

The experiment comprised four conditions: vestibular, auditory, and two combined-cue conditions (Δ = +6° and Δ = -6°). The task for all conditions was the same—to indicate whether the self-motion stimulus heading was to the left/right of straight ahead. Trials were initiated by pressing a central “start” button on the response box (Cedrus RB-540). Choices were reported after the stimulus had ended by pressing the corresponding right or left button on the response box. Three different auditory tones (unrelated to the self-motion stimuli), delivered via external speakers, were used to convey trial-timing information, as follows: (i) the system was ready for a new trial (i.e., to press start), (ii) a choice was registered, and (iii) a response time-out (if a choice was not registered within 2 s after the stimulus had ended). Participants were instructed to avoid this time-out by making a timely response, and to guess when unsure. No feedback was given regarding whether the choices were correct or incorrect. Before starting the experiment, the participants confirmed that they could hear these auditory tones from the external speakers (through the headphones) and were given a few practice trials to confirm that they understood the instructions well and performed the task reliably.

After each vestibular stimulus, the motion platform was required to return to its origin before beginning another trial. This return motion followed the reverse dynamics of the forward motion (along a straight line) however, it was performed more slowly (over 2.5 s vs. 1 s for the forward motion). To make the experience of the synthetic auditory cues more consistent and immersive, an auditory signal encoding the return motion was also played (on trials with auditory and combined-cue stimuli).

The four conditions were pseudo-randomly interleaved. On each trial, the heading sign was selected randomly (*p* = 0.5 for right or left). Task difficulty (heading magnitude) followed an adaptive staircase procedure (Cornsweet, 1962). A separate staircase was used per condition. Each staircase began with the easiest heading (±16°). After a correct choice, the heading magnitude was reduced (such that the task became more difficult) with *p* = 0.3, otherwise, it remained unchanged. After an incorrect response, the heading magnitude was increased (such that the task became easier) with *p* = 0.8, otherwise it remained unchanged. This staircase rule converges to ∼73% correct choices (Leek, 2001; Zaidel et al., 2015), thereby sampling an information-rich region of the psychometric function on an individual basis. A total of 400 trials were collected for each participant (100 trials per condition). This was divided into two blocks of 200 trials (lasting ∼20 min each) to allow for a break in the middle.

### Data analyses

Data analyses were performed using custom scripts in Matlab R2018b (The MathWorks). Psychometric functions were constructed, per condition, by calculating the proportion of rightward choices as a function of heading and fitting the data with a cumulative Gaussian distribution using the psignifit toolbox for Matlab, version 4 (Schütt et al., 2016). The psychometric fits had three free parameters: μ, σ, and λ. The first two parameters reflect the mean (μ) and SD (σ) of the cumulative Gaussian distribution function. These define the point of subjective equality (PSE or bias) and the psychophysical threshold, respectively. The third parameter (λ) reflects the lapse rate, which is the rate of reporting an incorrect choice for obvious stimuli. The lapse rates were assumed to be symmetrical for rightward and leftward headings. The three parameters were concurrently fit using the default toolbox priors. The goodness-of-fit of the psychometric functions was evaluated using pseudo-R² (Hosmer et al., 2013). Participants with pseudo-R² values lower than 0.5 for one of their psychometric functions (vestibular, auditory, or the two combined-cue conditions) were removed from further analysis. This excluded one participant. The pseudo-R² values for the remaining psychometric functions were: mean ± SD = 0.87 ± 0.07 for vestibular, 0.85 ± 0.13 for auditory, 0.88 ± 0.07 and 0.90 ± 0.07 for the combined-cue conditions (Δ = +6° and -6° respectively).

### Standard cue integration measures

The multisensory integration analyses were performed using the classic Bayesian equations, which have been well-described previously (Angelaki et al., 2009). Below, we briefly summarize the specific measurements used in this study (calculated per participant):

#### 1. *Observed* biases

The observed (empirical) vestibular and auditory biases (*μ*_*ves*_ and *μ*_*aud*_ respectively) were extracted from the unisensory psychometric fits.

#### 2. *Observed* thresholds

The observed (empirical) vestibular and auditory thresholds (*σ*_*ves*_and *σ*_*aud*_, respectively) were extracted from the unisensory psychometric fits. The observed combined-cue threshold (*σ*_*combined*_) was calculated by the (geometric) mean of the thresholds from the two combined-cue psychometric fits (from conditions Δ = +6° and -6°).

#### 3. Cue reliabilities

The vestibular and auditory cue reliabilities were estimated from the observed thresholds, as follows:

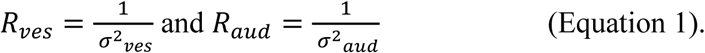

#### 4. Bayesian-predicted thresholds

The combined-cue thresholds were predicted, per participant, from their unisensory thresholds estimates (Ma et al., 2006), as follows:

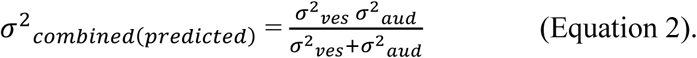

#### 5. Cue weights

##### 5.1. Bayesian-predicted cue weights

The Bayesian-predicted cue weights were calculated by the relative cue reliabilities, as follows:

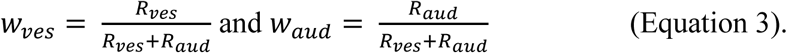

These weights sum to 1 (*w*_*aud*_ + *w*_*ves*_ = 1).

##### 5.2. *Observed* cue weights

The observed (empirical) auditory weights were estimated from the combined-cue conditions (Fetsch et al., 2009), as follows:

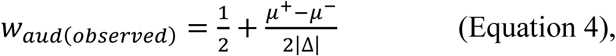

where *μ*^+^ and *μ*^−^ are the biases from the combined-cue conditions with positive and negative Δ, respectively. The corresponding observed vestibular weights are given by: *w*_*ves*(*observed*)_ = 1 − **w**_*aud*(*observed*)_.

### Combined-cue simulations

#### 1. Integration

Because combined-cue performance in this study (results presented below) was not in accordance with Bayesian-optimal predictions, we considered other possible models of cue combination. First, we performed a simulation of cue integration – similar to Bayesian integration – but allowing for a range of (non-optimal) weights. For simplicity, the unisensory vestibular and auditory thresholds were fixed: *σ*_*ves*_= 6° and *σ*_*aud*_ = 4° (these values reflect typical performance in the experiment).

Psychometric curves with different cue-integration weights were simulated using Δ = +6° (Fig. 2, A-C; Δ = -6° would give symmetrical results). For each cue, 10,000 trials were run per heading (ℎ = ±16°, ±8°, ±4°, ±2°, ±1°, ±0.5°, and ±0.25°). For unisensory cues, on each trial, a noisy heading estimate was drawn from a normal distribution around the stimulus heading (with offset) according to the respective cue’s threshold: *x*_*aud*_ ∼ *N* (ℎ − 3, *σ*^2^_*aud*_) or *x*_*ves*_∼ *N* (ℎ + 3, *σ*^2^_*ves*_). The simulated response (left or right) was then determined by comparing the noisy heading estimate to zero (a rightward choice for positive estimates and a leftward choice for negative estimates). For combined cues, a linear weighted sum of the unisensory estimates was compared to zero. The simulated data were fitted with psychometric curves (like the real data).

**Figure 2.**
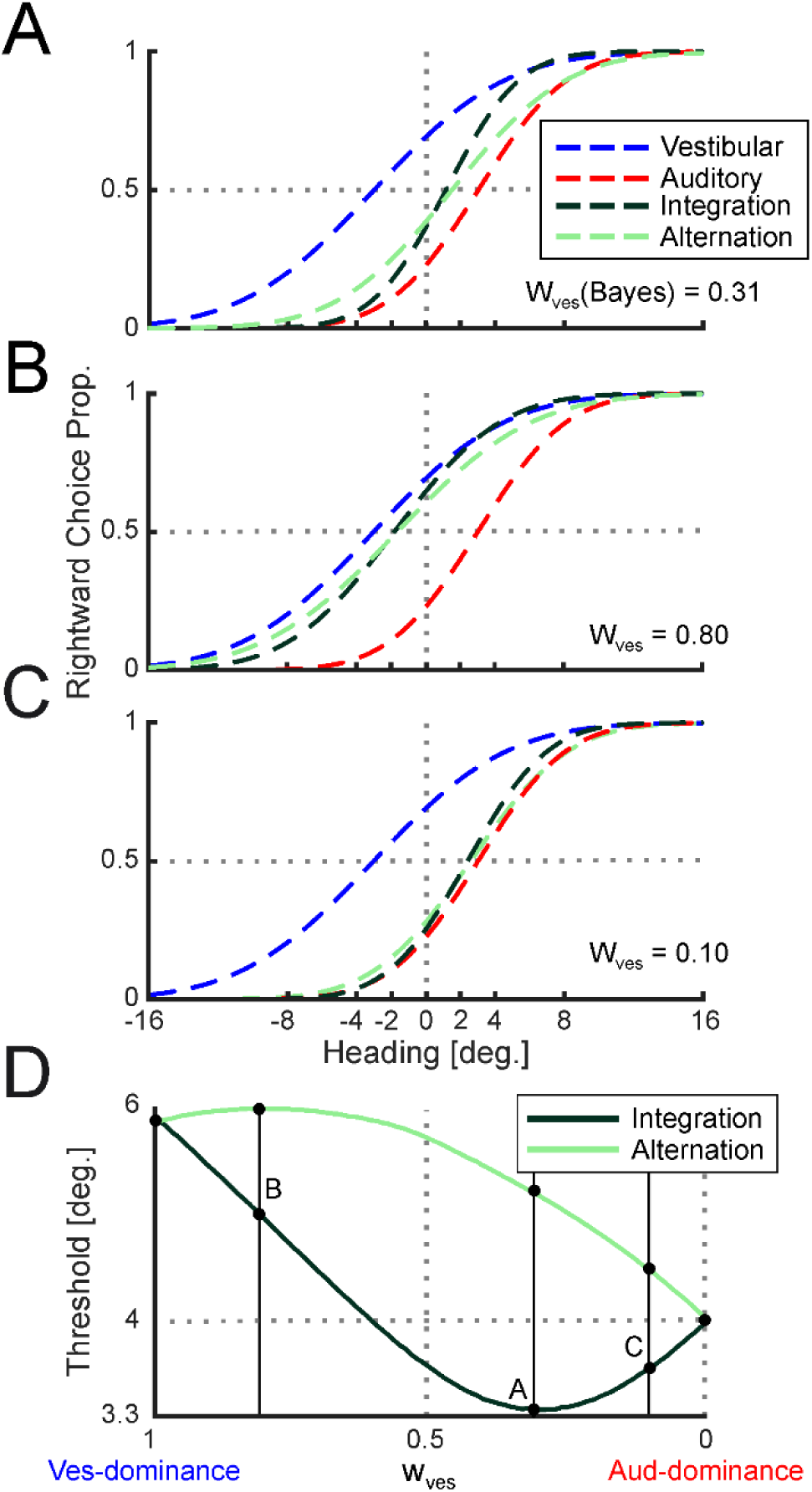
Simulated cue integration and cue alternation. (A-C) Psychometric curves (dashed lines) depict the simulated proportion of rightward choices as a function of heading for vestibular (blue) auditory (red) and two models of combined cues (*integration* and *alternation*, dark and light green, respectively). The vestibular and auditory curves were simulated using thresholds σ_ves_ = 6° and σ_aud_ = 4°, respectively. A heading discrepancy between the vestibular and auditory cues (Δ = 6°) was simulated by systematically shifting their headings +3° and -3°, respectively. Panels A-C present the same vestibular and auditory curves, but different combined curves, simulated using: (A) Bayesian-optimal weighting, (B) vestibular overweighting, and (C) vestibular underweighting. The vestibular weights used for the simulations are presented on the respective subplots (w_ves_; the corresponding auditory weights are given by: w_aud_ = 1 - w_ves_). (D) Combined-cue thresholds were simulated using the integration and alternation models across the full range of possible weights. The solid black vertical lines correspond to the simulations in panels A-C (as marked on the figure).

To assess combined-cue thresholds across the full range of possible integration weights (Fig. 2D) combined-cue curves were simulated using vestibular weights ranging from 0 to 1, in increments of 0.01 (with complementary auditory weights). This was performed with Δ = 0° because a heading discrepancy is not needed to estimate combined-cue thresholds. The simulation and psychometric fit were performed 100 times for each weight, and the thresholds were averaged using the geometric mean.

#### 2. Alternation

Besides cue integration, another potential model for combining cues is cue alternation (Goeke et al., 2016). For cue alternation, the weights represent the probability of selecting one unisensory cue’s estimate over the other. In this case, the combined-cue measurement on a given trial is equal to that of the selected cue alone (ignoring the other cue), unlike the weighted sum of cues for integration.

Psychometric curves with different cue-alternation weights (probabilities) were simulated using the same unisensory thresholds as the cue-integration simulation (Fig. 2, A-C). For the unisensory cues, noisy heading estimates were drawn in the same way as the cue-integration simulation. For combined cues, on each trial only one cue’s estimate (either vestibular or auditory) was selected, with probability according to the cues’ weights. This was considered the combined-cue estimate and compared to zero.

To assess combined-cue thresholds across the full range of possible alternation weights (Fig. 2D), combined-cue curves were simulated using vestibular weights ranging from 0 to 1, in increments of 0.01 with complementary auditory weights (similar to the integration simulation). In the alternation model (unlike the integration model) combined-cue threshold estimates are affected by the systematic heading discrepancy. Therefore, the alternation simulation was performed under two conditions: Δ = +6° and Δ = -6°. The simulated combined-cue threshold was calculated as the geometric mean of the thresholds obtained from the two psychometric curve fits. The simulation and psychometric fits were performed 100 times for each weight, and the thresholds were averaged using the geometric mean.

### Individual combined-cue threshold predictions

Using similar methods to the general simulations (described in the previous section, with fixed unisensory thresholds) we next simulated combined-cue threshold predictions for each participant, using their individual performance. This was done for both models (integration and alternation), using two different sets of weights: (i) Bayesian weights – predicted, per participant, from their unisensory thresholds (Eq. 3), and (ii) empirically observed weights – extracted from their combined-cue curves (Eq. 4). Like the general simulations, the integration model was simulated without a cue discrepancy (Δ = 0°), as this did not affect combined-cue thresholds. For the alternation model, combined-cue conditions were simulated using both discrepancies, as per the actual experiment (Δ = ±6°), and using each individual’s unisensory biases and thresholds.

### Statistical analysis

Statistical analyses were performed using JASP (version 0.19; JASP Team, 2024). To compare observed thresholds across the three cues (vestibular, auditory, and combined), a one-way repeated-measures ANOVA was used, with Bonferroni correction for post hoc pairwise comparisons. Sphericity was assessed using Mauchly’s test, and Greenhouse-Geisser corrections were applied when sphericity was violated. The pairwise comparison between vestibular and combined-cue thresholds (Fig 3C) was of particular interest, as it allowed us to assess whether participants used the synthetic auditory cue to improve perception. All statistical comparisons with thresholds used their logarithmic values (natural log), because thresholds are non-negative and scale logarithmically.

**Figure 3.**
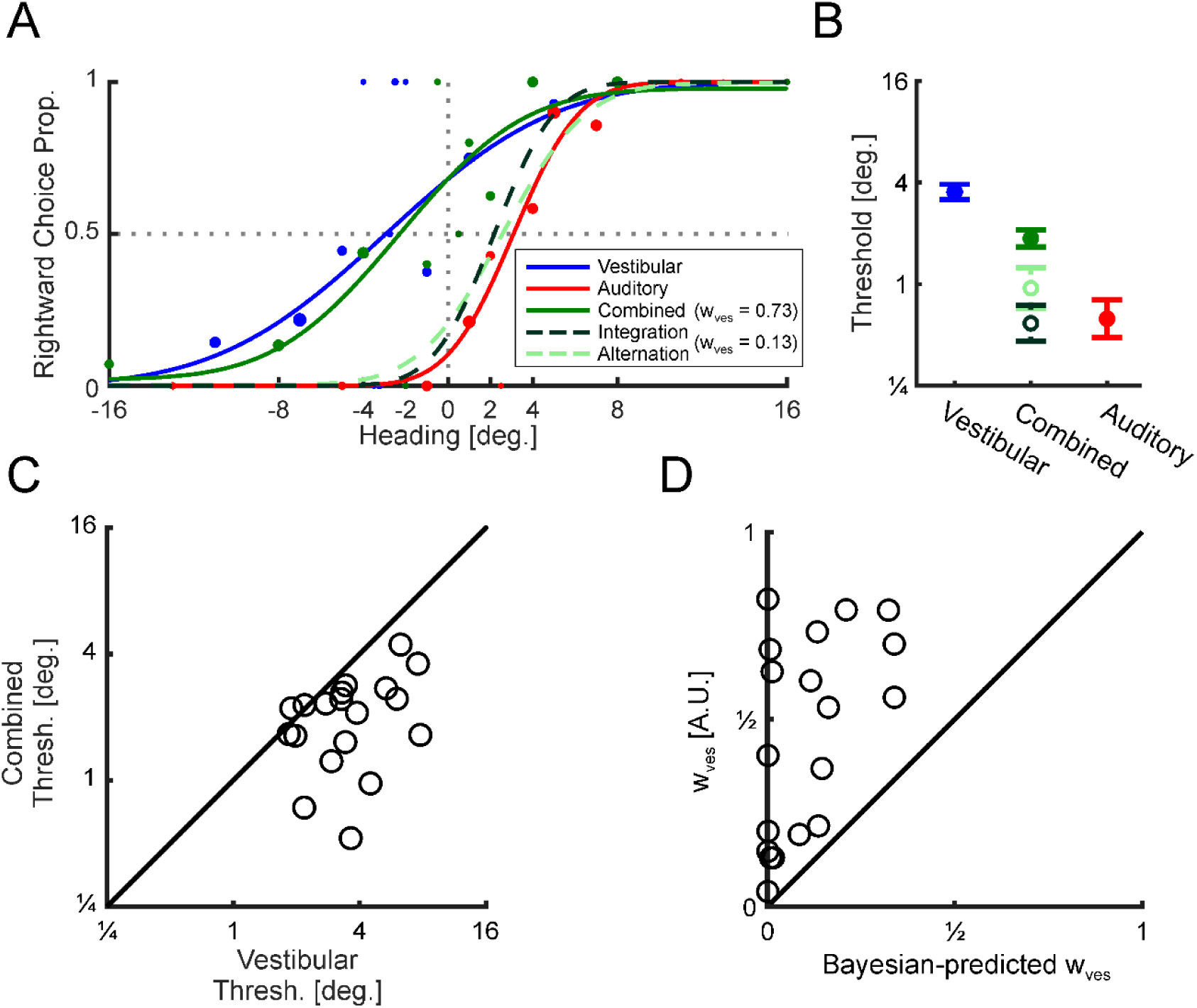
Synthetic auditory cues improve performance. (A) Psychometric data for an example participant. Datapoints mark the proportion of rightward choices as a function of heading for vestibular (blue), auditory (red), and combined (green) cues. Datapoint sizes reflect the number of trials collected at that heading. The data were fit, per cue, with psychometric curves (solid lines). For this example, the combined data were gathered with heading offset Δ = +6°. To reflect the offset, the vestibular and auditory responses were shifted (for presentation only) by 3° to the left and right, respectively. The dashed lines reflect simulated combine-cue curves for cue integration (dark green) and cue alternation (light green) using Bayesian-optimal weighting (w_ves_ = 0.13). By contrast, w_ves_ = 0.73 reflects the observed weighting (deduced from the actual combined-cue performance). The gray vertical dotted line marks heading = 0°, and the gray horizontal dotted line marks y = 0.5 (equal probability of rightward and leftward choices). (B) Filled circles mark the mean (± SEM) thresholds observed for vestibular, auditory, and combined cues. Open circles mark the mean (± SEM) simulated combined-cue thresholds for cue integration and cue alternation (dark and light green, respectively). (C) Observed thresholds: combined vs vestibular. (D) Observed vestibular weights (deduced from combined-cue performance) vs. those predicted by Bayesian theory (based on the vestibular and auditory cue reliabilities). The diagonal black lines in C and D mark the line of equality (y = x).

A two-tailed paired *t*-test and a Bayesian paired-sample *t*-test were used to compare the observed vestibular weights to the Bayesian-predicted vestibular weights (Fig. 3D). A similar comparison for auditory weights was not necessary because vestibular and auditory weights are complementary (sum to 1). Bayes factors (BF_10_) were interpreted as follows: BF_10_ > 3 indicates substantial evidence in favor of the alternative hypothesis (H_1_, that the observed weights differ from the Bayesian predictions), and BF_10_ > 10 suggests strong evidence for H_1_. Conversely, BF_10_ < 1/3 and BF_10_ < 1/10 provide substantial and strong evidence, respectively, for the null hypothesis (H_0_, that the observed weights are similar to the Bayesian predictions) (Jarosz & Wiley, 2014; Raftery, 1995; Wagenmakers, 2007).

To quantify (dis)similarity between the observed and the predicted combined-cue thresholds for the different models (Fig. 4) the distance of the data from the unity line (y = x) was measured using the normalized mean squared error (NMSE; normalized by the variance of the observations). To estimate variability in NMSE for each subplot in Figure 4, we performed bootstrap resampling with replacement (10,000 iterations) on the subject-level pairs of observed and model-predicted thresholds corresponding to each subplot. Because the distribution of NMSE values across bootstrap samples was right-skewed, we applied a log transformation before calculating summary statistics (mean and 95% confidence interval computed on the log scale and back-transformed for interpretability). For statistical comparisons, we computed the log-difference in NMSE between models using Bayesian versus observed weights (Fig. 4A vs. B) for both integration and alternation models, as well as between the integration and alternation models using observed weights (Fig. 4B, top vs. bottom subplots). Two-tailed p-values were obtained by calculating the proportion of bootstrap samples on each side of zero, taking the smaller of the two, and multiplying it by two.

**Figure 4.**
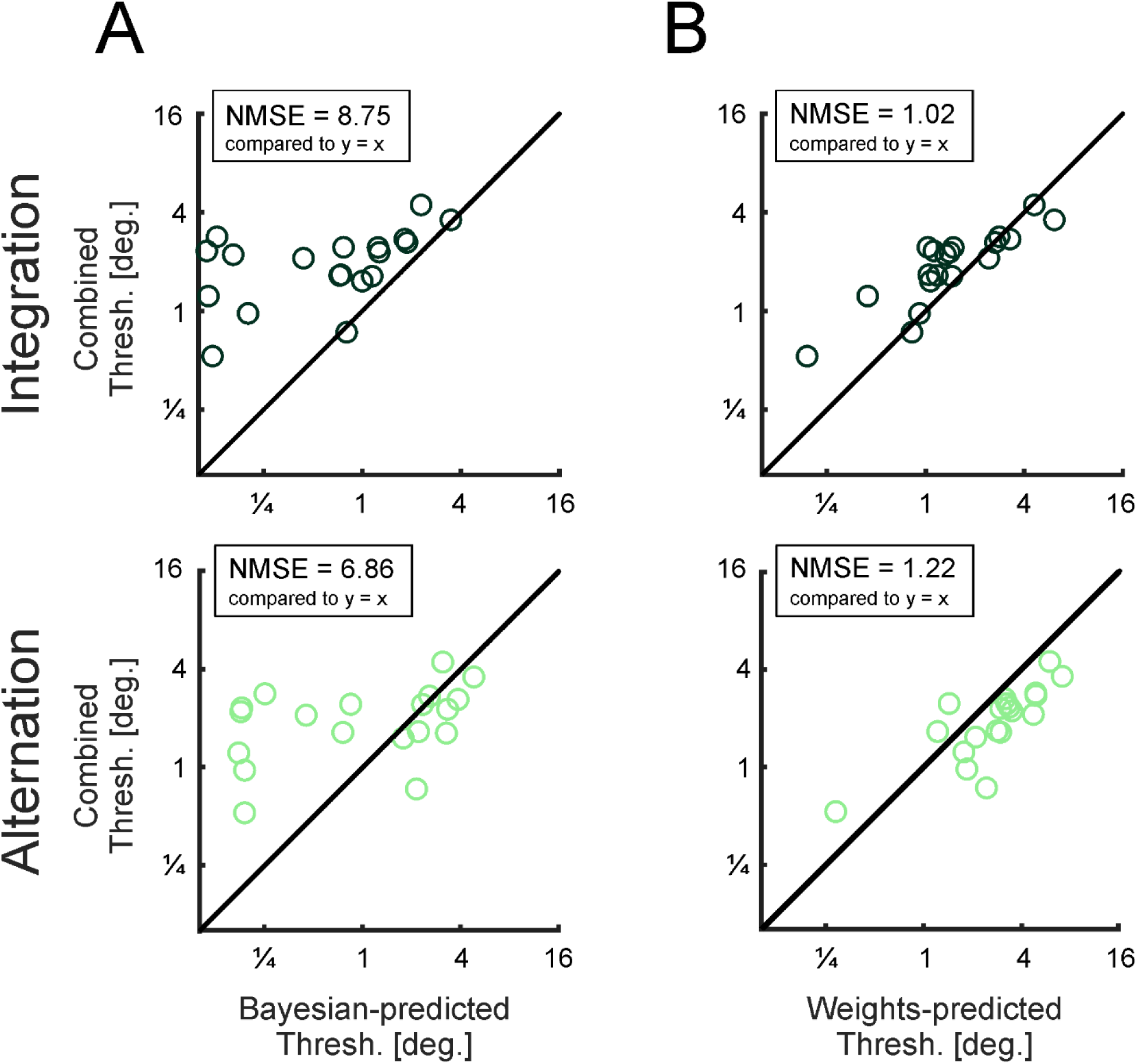
Combined-cue thresholds are better predicted using the observed vs. Bayesian weights. Scatter plots of the empirically observed combined-cue thresholds vs. model predictions using: Bayesian weights (A, left column) or the empirically observed weights (B, right column); and using a cue integration (top row) or cue alternation (bottom row) model. The normalized mean squared error (NMSE) was calculated per plot in reference to perfect prediction (y = x; marked by the diagonal black lines). Lower NMSE values mean better combined-cue threshold predictions.

### Code availability

The auditory stimulus files, data, and the analysis code for this study are available at: https://osf.io/qpwkg/?view_only=a5a39787d7424ab596e740ed0d58c9f6

## Results

### Cue integration and cue alternation simulation results

To demonstrate how combined-cue psychometric performance would manifest for non-optimal vs. optimal weights, we simulated three specific cases of interest: Bayesian-optimal weighting (Fig. 2A), vestibular overweighting (Fig. 2B), and vestibular underweighting (Fig. 2C). Moreover, this was done for two models/ strategies of combined-cue performance: cue integration and cue alternation. For all these cases, psychometric curves were simulated using *σ*_*aud*_ = 4° and *σ*_*ves*_ = 6° for auditory and vestibular thresholds, respectively (lower thresholds reflect better performance) and Δ = +6° for combined cues.

For Bayesian-optimal weighting (Fig. 2A) the vestibular cue was weighted less than the auditory cue, in accordance with their relative reliabilities (*w*_*ves*_= 0.31, *w*_*aud*_= 0.69; Eq. 3). In this case, the combined-cue curves (dark and light green, for integration and alternation, respectively) lie closer to the auditory (red) vs. vestibular (blue) curves. Thus, in terms of bias, cue integration and cue alternation were qualitatively similar. However, in terms of thresholds, they were quite different. For integration, the combined-cue threshold (*σ*_*combined*_ = 3.30°) was lower than both auditory and vestibular thresholds. Whereas, for alternation, the combined-cue threshold (*σ*_*combined*_ = 5.16°) was in between the two. Thus, while cue integration improved performance compared to unisensory cues, cue alternation improved performance only compared to the vestibular (but not the auditory) cue.

When the vestibular cue was overweighted (it was weighted more than the auditory cue, despite being less reliable: *w*_*ves*_ = 0.80, *w*_*aud*_= 0.20), the combined-cue curves (for both integration and alternation) were now closer to the vestibular curve (Fig. 2B). Overweighting the less-reliable cue increased the combined-cue thresholds for both cue integration and cue alternation, compared to the use of Bayesian-optimal weights (above). For cue integration, the combined-cue threshold (*σ*_*combined*_ = 4.86°) was still lower than the vestibular threshold. However, it was now higher than the auditory threshold. For cue alternation, the combined-cue threshold (*σ*_*combined*_ = 6.12°) was higher than both auditory and vestibular thresholds.

When underweighting the vestibular cue (*w*_*ves*_ = 0.10, *w*_*aud*_= 0.90) compared to the Bayesian prediction, the combined-cue curves lay close to the auditory curve (Fig. 2C). For cue integration, the combined-cue threshold (*σ*_*combined*_ = 3.67°) was lower than both the auditory and vestibular thresholds (but not as good as Bayesian-optimal integration). For cue alternation, the combined-cue threshold (*σ*_*combined*_ = 4.42°) was in between – lower than the vestibular threshold, but higher than the auditory threshold.

Beyond these three examples, we compared the simulated combined-cue thresholds, for both integration and alternation models, across the full range of weights (Fig. 2D). Overall, thresholds were higher for alternation (light green) compared to integration (dark green). For alternation, the lowest threshold occurred when relying solely on the more reliable (auditory) cue. Namely, including the less-reliable (vestibular) cue to any extent always yielded a worse threshold. Moreover, the combined-cue threshold could even be worse than the less reliable cue alone (e.g., Fig. 2B). By contrast, integration always yielded a better threshold than the less reliable (vestibular) cue. And, depending on the specific weights, the combined threshold could be higher, or lower, than the more reliable cue.

### Improved performance with a synthetic auditory cue

We next assess behavioral performance in the actual experiment. Psychometric curves of an example participant are presented in Figure 3A. In this example, we present the data from the combined-cue condition (solid green curve) with heading offset Δ = +6°. To reflect this offset, the vestibular and auditory responses (blue and red curves, respectively) were shifted (for presentation only) by 3° to the left and right, respectively. Performance was best for the auditory cue (steepest curve), worst for the vestibular cue, and in between for the combined-cue. This suggests that information from the synthetic auditory cue was indeed used in the combined-cue condition to improve performance.

At the group level, thresholds differed significantly across conditions (vestibular, auditory, and combined; *F*(1.42, 25.47) = 32.10, *p* = 9.00 × 10^-7^, *η*² = 0.64, repeated-measures ANOVA with Greenhouse-Geisser correction; Fig. 3B). In particular, combined-cue thresholds (2.10° ± 0.95°) were significantly lower than vestibular thresholds (3.90° ± 1.86°; *p* = 4.67 × 10^-4^, *t* _(18)_ = 4.76, Cohen’s *d* = 0.80; post hoc comparison with Bonferroni correction). This indicates that the presence of the auditory cue in the combined condition improved performance compared to vestibular alone. A scatter plot of each participant’s combined-cue vs vestibular thresholds (Fig. 3C) demonstrates improved combined-cue performance in most participants (17 out of 19 data points lie below the diagonal). Hence, the participants indeed used the synthetic auditory cue to improve self-motion perception. Additional post hoc comparisons showed that auditory thresholds (1.07° ± 1.01°) were significantly lower than vestibular thresholds (*p* = 9.98 × 10^-6^, *t* _(18)_ = 6.61, Cohen’s *d* = 2.18) and combined-cue thresholds (*p* = 6.76 × 10^-4^, *t* _(18)_ = 4.59, Cohen’s *d* = 1.38). Thus, auditory cues were more reliable than vestibular cues and, perhaps surprisingly, also more reliable than combined cues. This is addressed further below.

### Vestibular cues are overweighted

To quantitatively test whether the auditory and vestibular cues were integrated in a near-optimal Bayesian manner, we compared the empirically observed weights (Eq. 4) to the Bayesian predictions (Eq. 3). For the example in Figure 3A, the combined-cue (solid green) psychometric curve lies close to the vestibular (blue) curve. Accordingly, the empirically observed weights were larger for the vestibular vs. auditory cue (*w*_*ves*_= 0.73, and *w*_*aud*_= 0.27). This contrasts with the Bayesian prediction for a smaller vestibular vs. auditory weight (*w*_*ves*_ = 0.13; *w*_*aud*_ = 0.87; simulated combined-cue curves with the Bayesian-optimal weights are presented for the integration and alternation models, using dark and light green dashed lines, respectively). Thus, this example manifests vestibular overweighting.

In a scatter plot of the empirically observed vs. Bayesian-predicted vestibular weights, all the data points lie above the diagonal (Fig. 3D). This indicates that the observed vestibular weights were consistently higher than predicted. Overall, the vestibular cue was significantly overweighted compared to Bayesian predictions (*t* _(18)_ = 6.51, *p* = 4.04 × 10^-6^; Cohen’s *d* = 1.49, *t*-test, BF_10_ = 5020). Hence, empirical cue weighting was not based solely on cue reliability.

Given the substantial vestibular overweighting, it is not surprising that the combined-cue thresholds were not well-predicted using Bayesian weights (neither with the integration nor alternation models; Fig. 3B). Demonstrating this at the individual level, scatter plots of the observed combined-cue thresholds vs. those predicted using Bayesian weights are shown in Figure 4A (top and bottom subplots for the integration and alternation models, respectively). The high NMSE values (presented on the subplots) indicate substantial discrepancies between the models’ predictions and the observed combined-cue thresholds (i.e., divergence of the data from the diagonal).

### Combined-cue thresholds are better-predicted by the empirically observed weights

Up to this point, we have shown that although the synthetic auditory cues did improve performance compared to vestibular cues alone, they were not integrated with the vestibular cues in a Bayesian (reliability-based) manner. This may not be surprising, given that the auditory cue was synthetic, not previously associated with self-motion, and thus likely considered less relevant for the task. Accordingly, we considered that the participants might have still performed cue integration (or alternation), but with weights that differ from the reliability-based predictions. Therefore, we tested whether cue integration (or cue alternation) using the empirically observed (rather than Bayesian) weights could better predict the combined cue thresholds.

Indeed, when using the empirical weights to predict the combined-cue thresholds, scatter plots of the observed vs. predicted thresholds lay more closely to the diagonal, for both the integration and alternation models (Fig. 4B, top and bottom subplots, respectively). Correspondingly, the NMSE values (presented on the subplots) were close to 1, and significantly lower vs. those with Bayesian-predicted weighting. NMSE = 1.02 [0.45, 2.57] for cue integration using the empirical weights vs. 8.75 [3.04, 27.77] using Bayesian weights (*p* = 3.80 × 10⁻³). NMSE = 1.22 [0.64, 2.60] for cue alternation using empirical weights vs. 6.86 [2.46, 21.41] using Bayesian weights (*p* = 9.20 × 10⁻³). Thus, for both models, the observed combined-cue thresholds were better predicted using empirical rather than reliability-based weights. The NMSE, when using empirical weights, was marginally (but not significantly) lower for integration vs. alternation (*p* = 0.72).

## Discussion

In this study, we investigated whether providing a synthetic auditory cue, designed to encode linear self-motion in the horizontal plane, could augment vestibular signals and thereby improve self-motion perception, in healthy participants. Heading discrimination thresholds were significantly better with combined auditory-vestibular cues than with the vestibular cue alone. Thus, the synthetic auditory cue indeed improved self-motion perception. However, combined-cue performance was not Bayesian-optimal. Specifically, vestibular cues were heavily overweighted compared to Bayesian (reliability-based) predictions. Moreover, combined-cue thresholds were worse than predicted by Bayesian-optimal integration. In fact, they were even worse than the unimodal auditory thresholds (which were surprisingly reliable).

Although slight vestibular overweighting was also noted in visual-vestibular heading discrimination (Butler et al., 2010; Fetsch et al., 2009), it was relatively subtle. There, combined-cue thresholds were lower vs. both visual or vestibular cues alone, reflecting near-optimal integration. The difference is likely explained by the fact that visual and vestibular cues are both naturally associated with self-motion, unlike the auditory cue used in this study, which was synthetic and unfamiliar to the participants. In line with this idea, near-optimal integration has been consistently observed in many other paradigms where the sensory cues are naturally suited to the task. For example, visual and haptic cues for size perception (Ernst & Banks, 2002), auditory and visual cues for object localization (Alais & Burr, 2004), and visual and proprioceptive cues for hand localization (van Beers et al., 1999).

Our findings here align with other recent studies demonstrating that while novel cues can be integrated with natural cues, the integration is suboptimal (Negen et al., 2023; Shayman et al., 2025). In studies by Negen et al. (2018, 2023), participants learned in just a few hours of training to use an echo-like synthetic auditory cue to judge the distance to a target. When the auditory cue was paired with vision, perceptual reliability improved, although not to the full extent predicted by Bayesian-optimal integration. Rather, participants tended to overweight the visual (native) cue and underweight the auditory (synthetic) cue. A similar pattern to underweight unfamiliar auditory cues was also recently found in a virtual reality homing task, where participants relied more heavily on body-based self-motion cues to navigate vs. spatial auditory landmarks (Shayman et al., 2025). Taken together, these findings and our results suggest that while novel synthetic cues can be integrated with natural cues to improve perceptual function, the benefit falls short of its full potential.

We consider two possible explanations for the underweighting of synthetic cues. One possibility is that participants may have judged their perception of the synthetic auditory cue to be less reliable than its objective (experimentally measured) statistical reliability. Indeed, metacognitive assessments of reliability can diverge from the actual reliability of task performance (Fleming, 2024). The very fact that the auditory cue was unfamiliar might have impacted the brain’s ability to accurately estimate its reliability. In this case, the integration itself could be in line with Bayesian principles – i.e., cue-weighing according to their (estimated) relative reliabilities – but using an undervalued estimate of the synthetic auditory cue’s reliability.

An alternative explanation is that perceptual reliability of the synthetic auditory cue was correctly estimated, but the brain also takes into account other sources of uncertainty, besides reliability (Aston et al., 2022). For instance, a synthetic cue may be down-weighted not because it is unreliable, but because it lacks prior association with the task and is therefore considered less relevant. This is part of the brain’s broader ‘causal inference’ problem: deciding which sensory signals should be integrated. According to Bayesian models of causal inference (Körding et al., 2007) the brain can partially integrate the cues – namely, form an overall estimate based on a probabilistic combination of both integration and segregation. In the case of a novel synthetic cue, the brain might be uncertain whether or not to integrate it with the natural cue. This might explain why we found vestibular overweighting. Because, in the context of causal inference, the final heading estimate could be a combination of the integrated (auditory-vestibular) and segregated (vestibular) measures. To test this idea, future research is needed with study designs that can differentiate causal inference from simple overweighting.

Causal inference for multisensory integration relies heavily on past experience (Zaidel & Salomon, 2023), and co-occurrence of multisensory signals during development plays a critical role in shaping such expectations (Stein et al., 2014; Wallace, 2004). When a novel cue is introduced, the brain may require time to adjust its internal model, to treat it as a relevant source of information that should be integrated. Thus, while the brain can rapidly adjust integration weights for cues that are naturally associated with the task – based on instantaneous estimates of their reliability (Fetsch et al., 2009) – updating the assessment of a cue’s relevance to the task may be a much slower process, that requires prolonged exposure or training (Daee et al., 2014; Weisswange et al., 2011). Indeed, training with novel sensory cues can improve integration and shift cue weighting toward Bayesian-optimal predictions (Aston et al., 2023; Negen et al., 2018).

Combined-cue thresholds were much better predicted when using the individuals’ empirically observed (vs. Bayesian-optimal) weights. This further confirms that they were indeed over-(under-) weighting their vestibular (auditory) cues in the combined cue conditions. Although a model of cue integration vs. cue alternation did not provide a significantly better fit for the combined cue thresholds, the error tended to be smaller for integration vs. alternation when using the empirical weights. Also, the combined-cue thresholds were consistently lower vs. the (less reliable) vestibular cue alone (for 17/19 participants). This pattern was more consistent with cue integration, as simulations of cue alternation for our combined-cue conditions (with Δ) provided thresholds that were larger than those for the vestibular cue alone, when it was overweighted (as illustrated in Fig. 2D, point B). Thus, our results are more consistent with integration using overweighted vestibular cues.

Although in this study we tested only healthy participants, its findings may have particular relevance for individuals with vestibular dysfunction. Such individuals have been found to rely more heavily on auditory cues for spatial orientation and postural control (Shayman et al., 2018; Stevens et al., 2017; Vitkovic et al., 2016). Thus, auditory cues can be used for self-motion perception when necessary (i.e., vestibular input is impaired) and individuals with vestibular dysfunction may already exhibit a predisposition to use auditory cues. This raises the possibility that auditory augmentation could benefit such individuals, provided they can, with training learn to use synthetic auditory cues together with their remaining vestibular function. One limitation of this study is that it tested only linear self-motions in the horizontal plane. In everyday life, self-motions involve additional dimensions, including vertical translations and rotational motions such as pitch, yaw, and roll. Further research is needed to determine whether synthetic auditory cues can improve perception in these dimensions, and which auditory features, such as pitch, volume, and timbre, are best suited to each dimension. Moreover, future research is needed to determine whether people can effectively use composite auditory signals that simultaneously encode multiple motion components, and what training would best achieve this goal.

Another limitation of this study is that it was performed with seated participants, in a controlled laboratory setting. To evaluate the practical utility of synthetic auditory cues for vestibular sensory augmentation, it will be essential to test function during active, self-generated movements, such as walking and balancing. Lastly, finding the best algorithm(s) for translating vestibular to auditory cues, to enhance learning and integration, is a fundamental challenge for future work on this topic

## Acknowledgments

We would like to thank Avraham Elkaras for programming assistance and Tamar Harpaz for administrative assistance. This work was supported by grants from the Israel Innovation Authority (Kamin) and from the Israel Science Foundation (ISF, grant No. 1291/20) to AZ.

## References

Acerbi, L., Dokka, K., Angelaki, D. E., & Ma, W. J. (2018). Bayesian comparison of explicit and implicit causal inference strategies in multisensory heading perception. PLoS Computational Biology, 14(7), e1006110. 10.1371/journal.pcbi.1006110

Alais, D., & Burr, D. (2004). The Ventriloquist Effect Results from Near-Optimal Bimodal Integration. Current Biology, 14(3), 257–262. 10.1016/J.CUB.2004.01.029

Allen, D., Ribeiro, L., Arshad, Q., & Seemungal, B. M. (2016). Age-Related Vestibular Loss: Current Understanding and Future Research Directions. Frontiers in Neurology, 7, 231. 10.3389/FNEUR.2016.00231

Angelaki, D. E., Gu, Y., & DeAngelis, G. C. (2009). Multisensory integration: Psychophysics, neurophysiology, and computation. Current Opinion in Neurobiology, 19(4), 452–458. 10.1016/j.conb.2009.06.008

Anson, E., & Jeka, J. (2016). Perspectives on aging vestibular function. Frontiers in Neurology, 6, 269. 10.3389/FNEUR.2015.00269

Aston, S., Beierholm, U., & Nardini, M. (2022). Newly Learned Novel Cues to Location Are Combined With Familiar Cues but Not Always With Each Other. Journal of Experimental Psychology: Human Perception and Performance, 48(6), 639–652. 10.1037/xhp0001014

Aston, S., Nardini, M., & Beierholm, U. (2023). Different types of uncertainty in multisensory perceptual decision making. Philosophical Transactions of the Royal Society B, 378(1886). 10.1098/RSTB.2022.0349

Auvray, M., Hanneton, S., & O’Regan, J. K. (2007). Learning to Perceive with a Visuo– Auditory Substitution System: Localisation and Object Recognition with ‘The Voice’. Perception, 36(3), 416–430. 10.1068/p5631

Bach-y-Rita, P., Collins, C. C., Saunders, F. A., White, B., & Scadden, L. (1969). Vision substitution by tactile image projection. Nature, 221(5184), 963–964. 10.1038/221963A0

Bach-y-Rita, P., Danilov, Y., Tyler, M. E., & Grimm, R. J. (2005). Late human brain plasticity: Vestibular substitution with a tongue BrainPort human-machine interface. Intellectica, 40(1), 115–122. 10.3406/intel.2005.1362

Bergen, G., Stevens, M. R., & Burns, E. R. (2016). Falls and fall injuries among adults aged ≥65 years—United States, 2014. Morbidity and Mortality Weekly Report, 65(37), 938–983. 10.15585/mmwr.mm6537a2

Boutros, P. J., Schoo, D. P., Rahman, M., Valentin, N. S., Chow, M. R., Ayiotis, A. I., Morris, B. J., Hofner, A., Rascon, A. M., Marx, A., Deas, R., Fridman, G. Y., Davidovics, N. S., Ward, B. K., Treviño, C., Bowditch, S. P., Roberts, D. C., Lane, K. E., Gimmon, Y., … Della Santina, C. C. (2019). Continuous vestibular implant stimulation partially restores eye-stabilizing reflexes. JCI Insight, 4(22), e128397. 10.1172/JCI.INSIGHT.128397

Burton, R. (2015). The elements of music: What are they, and who cares? Music: Educating for Life – Proceedings of the ASME XXth National Conference, 22–28. https://search.informit.org/doi/10.3316/informit.649996699786780

Butler, J. S., Smith, S. T., Campos, J. L., & Bülthoff, H. H. (2010). Bayesian integration of visual and vestibular signals for heading. Journal of Vision, 10(11), 23. 10.1167/10.11.23

Chow, M. R., Ayiotis, A. I., Schoo, D. P., Gimmon, Y., Lane, K. E., Morris, B. J., Rahman, M. A., Valentin, N. S., Boutros, P. J., Bowditch, S. P., Ward, B. K., Sun, D. Q., Treviño Guajardo, C., Schubert, M. C., Carey, J. P., & Della Santina, C. C. (2021). Posture, Gait, Quality of Life, and Hearing with a Vestibular Implant. New England Journal of Medicine, 384(6), 521–532. 10.1056/NEJMOA2020457

Cornsweet, T. N. (1962). The staircase-method in psychophysics. The American Journal of Psychology, 75, 485–491. 10.2307/1419876

Daee, P., Mirian, M. S., & Ahmadabadi, M. N. (2014). Reward Maximization Justifies the Transition from Sensory Selection at Childhood to Sensory Integration at Adulthood. PLoS ONE, 9(7), e103143. 10.1371/journal.pone.0103143

De Winkel, K. N., Katliar, M., & Bülthoff, H. H. (2017). Causal inference in multisensory heading estimation. PLoS ONE, 12(1), e0169676. 10.1371/journal.pone.0169676

Dozza, M., Chiari, L., & Horak, F. B. (2005). Audio-biofeedback improves balance in patients with bilateral vestibular loss. Archives of Physical Medicine and Rehabilitation, 86(7), 1401–1403. 10.1016/j.apmr.2004.12.036

Ernst, M. O., & Banks, M. S. (2002). Humans integrate visual and haptic information in a statistically optimal fashion. Nature, 415(6870), 429–433. 10.1038/415429A

Fetsch, C. R., Turner, A. H., DeAngelis, G. C., & Angelaki, D. E. (2009). Dynamic reweighting of visual and vestibular cues during self-motion perception. Journal of Neuroscience, 29(49), 15601–15612. 10.1523/JNEUROSCI.2574-09.2009

Fleming, S. M. (2024). Metacognition and Confidence: A Review and Synthesis. Annual Review of Psychology, 75, 241–268. 10.1146/annurev-psych-022423-032425

Florence, C. S., Bergen, G., Atherly, A., Burns, E., Stevens, J., & Drake, C. (2018). Medical Costs of Fatal and Nonfatal Falls in Older Adults. Journal of the American Geriatrics Society, 66(4), 693–698. 10.1111/jgs.15304

Fortin, M., Voss, P., Lord, C., Lassonde, M., Pruessner, J., Saint-Amour, D., Rainville, C., & Lepore, F. (2008). Wayfinding in the blind: Larger hippocampal volume and supranormal spatial navigation. Brain, 131(11), 2995–3005. 10.1093/brain/awn250

Gandemer, L., Parseihian, G., Kronland-Martinet, R., & Bourdin, C. (2017). Spatial cues provided by sound improve postural stabilization: Evidence of a spatial auditory map? Frontiers in Neuroscience, 11, 357. 10.3389/FNINS.2017.00357

Goeke, C. M., Planera, S., Finger, H., & König, P. (2016). Bayesian alternation during tactile augmentation. Frontiers in Behavioral Neuroscience, 10, 211879. 10.3389/FNBEH.2016.00187

Golub, J. S., Ling, L., Nie, K., Nowack, A., Shepherd, S. J., Bierer, S. M., Jameyson, E., Kaneko, C. R. S., Phillips, J. O., & Rubinstein, J. T. (2014). Prosthetic implantation of the human vestibular system. Otology and Neurotology, 35(1), 136–147. 10.1097/MAO.0000000000000003

Gu, Y., DeAngelis, G. C., & Angelaki, D. E. (2007). A functional link between area MSTd and heading perception based on vestibular signals. Nature Neuroscience, 10(8), 1038–1047. 10.1038/nn1935

Guinand, N., Van De Berg, R., Ranieri, M., Cavuscens, S., Digiovanna, J., Nguyen, T. A. K., Micera, S., Stokroos, R., Kingma, H., Guyot, J. P., & Perez Fornos, A. (2015). Vestibular implants: Hope for improving the quality of life of patients with bilateral vestibular loss. Proceedings of the 2015 Annual International Conference of the IEEE Engineering in Medicine and Biology Society (EMBC), 7192–7195. 10.1109/embc.2015.7320051

Hasegawa, N., Takeda, K., Sakuma, M., Mani, H., Maejima, H., & Asaka, T. (2017). Learning effects of dynamic postural control by auditory biofeedback versus visual biofeedback training. Gait & Posture, 58, 188–193. 10.1016/J.GAITPOST.2017.08.001

Hegeman, J., Honegger, F., Kupper, M., & Allum, J. H. J. (2005). The balance control of bilateral peripheral vestibular loss subjects and its improvement with auditory prosthetic feedback. Journal of Vestibular Research, 15(2), 109–117. 10.3233/VES-2005-15206

Hosmer, D. W., Lemeshow, S., & Sturdivant, R. X. (2013). Applied Logistic Regression: Third Edition. Applied Logistic Regression: Third Edition, 1–510. 10.1002/9781118548387

Huffman, J. L., Norton, L. E., Adkin, A. L., & Allum, J. H. J. (2010). Directional effects of biofeedback on trunk sway during stance tasks in healthy young adults. Gait & Posture, 32(1), 62–66. 10.1016/j.gaitpost.2010.03.009

Iwasaki, S., & Yamasoba, T. (2015). Dizziness and imbalance in the elderly: Age-related decline in the vestibular system. Aging and Disease, 6(1), 38–47. 10.14336/ad.2014.0128

Jacobson, G. P., McCaslin, D. L., Grantham, S. L., & Piker, E. G. (2008). Significant vestibular system impairment is common in a cohort of elderly patients referred for assessment of falls risk. Journal of the American Academy of Audiology, 19(10), 799–807. 10.3766/jaaa.19.10.7

Jarosz, A. F., & Wiley, J. (2014). What Are the Odds? A Practical Guide to Computing and Reporting Bayes Factors. The Journal of Problem Solving, 7(1), 2. 10.7771/1932-6246.1167

*JASP (Version 0.19) [Computer software]* (Version 0.19). (2024). [Computer software]. JASP Team. https://jasp-stats.org/

Körding, K. P., Beierholm, U., Ma, W. J., Quartz, S., Tenenbaum, J. B., & Shams, L. (2007). Causal inference in multisensory perception. PLoS ONE, 2(9). 10.1371/journal.pone.0000943

Leek, M. R. (2001). Adaptive procedures in psychophysical research. Perception and Psychophysics, 63(8), 1279–1292. 10.3758/BF03194543

Lopez, C., Blanke, O., & Mast, F. W. (2012). The human vestibular cortex revealed by coordinate-based activation likelihood estimation meta-analysis. Neuroscience, 212, 159–179. 10.1016/j.neuroscience.2012.03.028

Ma, W. J., Beck, J. M., Latham, P. E., & Pouget, A. (2006). Bayesian inference with probabilistic population codes. Nature Neuroscience, 9(11), 1432–1438. 10.1038/nn1790

Mahmud, M. R., Stewart, M., Cordova, A., & Quarles, J. (2022). Auditory Feedback for Standing Balance Improvement in Virtual Reality. Proceedings of the 2022 IEEE Conference on Virtual Reality and 3D User Interfaces (VR), 782–791. 10.1109/vr51125.2022.00100

Maidenbaum, S., Abboud, S., & Amedi, A. (2014). Sensory substitution: Closing the gap between basic research and widespread practical visual rehabilitation. Neuroscience and Biobehavioral Reviews, 41, 3–15. 10.1016/j.neubiorev.2013.11.007

Negahban, H., Bavarsad Cheshmeh ali, M., & Nassadj, G. (2017). Effect of hearing aids on static balance function in elderly with hearing loss. Gait and Posture, 58, 126–129. 10.1016/j.gaitpost.2017.07.112

Negen, J., Bird, L.-A., Slater, H., Thaler, L., & Nardini, M. (2023). Multisensory Perception and Decision-Making With a New Sensory Skill. Journal of Experimental Psychology: Human Perception and Performance, 49(5), 600–622. 10.1037/xhp0001114

Negen, J., Wen, L., Thaler, L., & Nardini, M. (2018). Bayes-Like Integration of a New Sensory Skill with Vision. Scientific Reports, 8(1), 1–12. 10.1038/s41598-018-35046-7

Raftery, A. E. (1995). Bayesian Model Selection in Social Research. Sociological Methodology, 25, 111–163. 10.2307/271063

Rancz, E. A., Moya, J., Drawitsch, F., Brichta, A. M., Canals, S., & Margrie, T. W. (2015). Widespread vestibular activation of the rodent cortex. Journal of Neuroscience, 35(15), 5926–5934. 10.1523/JNEUROSCI.1869-14.2015

Schütt, H. H., Harmeling, S., Macke, J. H., & Wichmann, F. A. (2016). Painfree and accurate Bayesian estimation of psychometric functions for (potentially) overdispersed data. Vision Research, 122, 105–123. 10.1016/J.VISRES.2016.02.002

Shalom-Sperber, S., Chen, A., & Zaidel, A. (2022). Rapid cross-sensory adaptation of self-motion perception. Cortex, 148, 14–30. 10.1016/j.cortex.2021.11.018

Shayman, C. S., Mancini, M., Weaver, T. S., King, L. A., & Hullar, T. E. (2018). The contribution of cochlear implants to postural stability. Laryngoscope, 128(7), 1676–1680. 10.1002/lary.26994

Shayman, C. S., Mccracken, M. K., Finney, H. C., Fino, P. C., Stefanucci, J. K., Creem-Regehr, S. H., Katsanevas, A., Carter, L., Gottschalk, H., Morse, M., Myers, M., Oldfield, L., Holm, G., & Schmidt, T. (2025). Integration of auditory and self-motion cues in spatial navigation. Journal of Experimental Psychology: Human Perception and Performance, 51(5), 664–675. 10.1037/xhp0001316

Sienko, K. H., Balkwill, M. D., Oddsson, L. I. E., & Wall, C. (2008). Effects of multi-directional vibrotactile feedback on vestibular-deficient postural performance during continuous multi-directional support surface perturbations. Journal of Vestibular Research, 18, 273–285. 10.3233/ves-2008-185-604

Sluydts, M., Curthoys, I., Vanspauwen, R., Papsin, B. C., Cushing, S. L., Ramos, A., Ramos De Miguel, A., Borkoski Barreiro, S., Barbara, M., Manrique, M., & Zarowski, A. (2020). Electrical Vestibular Stimulation in Humans: A Narrative Review. Audiology and Neurotology, 25(1–2), 6–24. 10.1159/000502407

Spagnol, S., Baldan, S., & Unnthorsson, R. (2017). Auditory depth map representations with a sensory substitution scheme based on synthetic fluid sounds. 2017 IEEE 19th International Workshop on Multimedia Signal Processing (MMSP), 1–6. 10.1109/MMSP.2017.8122220

Stein, B. E., Stanford, T. R., & Rowland, B. A. (2014). Development of multisensory integration from the perspective of the individual neuron. Nature Reviews Neuroscience, 15(8), 520–537. 10.1038/nrn3742

Stevens, M. N., Barbour, D. L., Gronski, M. P., & Hullar, T. E. (2017). Auditory contributions to maintaining balance. Journal of Vestibular Research: Equilibrium and Orientation, 26(5– 6), 433–438. 10.3233/VES-160599

van Beers, R. J., Sittig, A. C., & Gon, J. J. (1999). Integration of proprioceptive and visual position-information: An experimentally supported model. Journal of Neurophysiology, 81(3), 1355–1364. 10.1152/jn.1999.81.3.1355

Velázquez, R. (2010). Wearable assistive devices for the blind. In Lecture Notes in Electrical Engineering (Vol. 75, pp. 331–349). Springer. 10.1007/978-3-642-15687-8_17

Vitkovic, J., Le, C., Lee, S. L., & Clark, R. A. (2016). The Contribution of Hearing and Hearing Loss to Balance Control. Audiology and Neurotology, 21(4), 195–202. 10.1159/000445100

Wagenmakers, E. J. (2007). A practical solution to the pervasive problems of p values. Psychonomic Bulletin & Review, 14(5), 779–804. 10.3758/BF03194105

Wallace, M. T. (2004). The development of multisensory processes. Cognitive Processing, 5(2), 69–83. 10.1007/S10339-004-0017-Z

Weisswange, T. H., Rothkopf, C. A., Rodemann, T., & Triesch, J. (2011). Bayesian Cue Integration as a Developmental Outcome of Reward Mediated Learning. PLOS ONE, 6(7), e21575. 10.1371/journal.pone.0021575

Yakubovich, S., Israeli-Korn, S., Halperin, O., Yahalom, G., Hassin-Baer, S., & Zaidel, A. (2020). Visual self-motion cues are impaired yet overweighted during visual–vestibular integration in Parkinson’s disease. Brain Communications, 2(1). 10.1093/BRAINCOMMS/FCAA035

Zaidel, A. (2024). Multisensory Calibration: A Variety of Slow and Fast Brain Processes Throughout the Lifespan. In Multisensory Processes Across the Lifespan (Vol. 1437, pp. 173–201). Springer. 10.1007/978-981-99-7611-9_9

Zaidel, A., Goin-Kochel, R. P., & Angelaki, D. E. (2015). Self-motion perception in autism is compromised by visual noise but integrated optimally across multiple senses. Proceedings of the National Academy of Sciences, 112(20), 6461–6466. 10.1073/PNAS.1506582112

Zaidel, A., Ma, W. J., & Angelaki, D. E. (2013). Supervised calibration relies on the multisensory percept. Neuron, 80(6), 1544–1557. 10.1016/J.NEURON.2013.09.026

Zaidel, A., & Salomon, R. (2023). Multisensory decisions from self to world. Philosophical Transactions of the Royal Society B: Biological Sciences, 378(1886), 20220335. 10.1098/rstb.2022.0335

Zaidel, A., Turner, A. H., & Angelaki, D. E. (2011). Multisensory calibration is independent of cue reliability. Journal of Neuroscience, 31(39), 13949–13962. 10.1523/jneurosci.2732-11.2011

Zeng, F., Zaidel, A., & Chen, A. (2023). Contrary neuronal recalibration in different multisensory cortical areas. eLife, 12, e82895. 10.7554/ELIFE.82895

